# Rapid identification of optimal drug combinations for personalized cancer therapy using microfluidics

**DOI:** 10.1101/093906

**Authors:** Federica Eduati, Ramesh Utharala, Dharanija Madhavan, Ulf Peter Neumann, Thorsten Cramer, Julio Saez-Rodriguez, Christoph A. Merten

**Author notes:** co-first authors (Federica Eduati, Ramesh Utharala).

## Abstract

Functional screening of live patient cancer cells holds great potential for personalized medicine and allows to overcome the limited translatability of results from existing in-vitro and ex-vivo screening models. Here we present a plug-based microfluidics approach enabling the testing of drug combinations directly on cancer cells from patient biopsies. The entire procedure takes less than 48 hours after surgery and does not require ex vivo cultivation. We screened more than 1100 samples for different primary human tumors (each with 56 conditions and at least 20 replicates), and obtained highly specific sensitivity profiles. This approach allowed us to derive optimal treatment options which we further validated in two different pancreatic cancer cell lines. This workflow should pave the way for rapid determination of optimal personalized cancer therapies at assay costs of less than US$ 150 per patient.

## Introduction

Increasing interest has been devoted to personalized (precision) medicine, which aims to match the best treatment to each patient. This is especially important for cancer, where there is often a high variability in the response of patients to treatments. A critical example is pancreatic cancer, which has a 5-year survival rate of only 6% and is projected to become the second leading cause of cancer death by 2030 ^1^. One of the reasons for the low efficacy of current treatments is that pancreatic cancer is genetically highly heterogeneous with most mutations occurring at a prevalence of less than 10% ^2,3^. Therefore, pancreatic cancer patients would benefit immensely from the possibility of a personalized or stratified therapeutic approach.

Most efforts in personalized medicine have been focusing on tailoring the treatment to the specific patient based on genomic data, which are increasingly available due to advances in sequencing technologies. While there have been some impressively successful examples ^4,5^, cancer genomics is generally very complex and, despite the increasing knowledge on occurring mutations, there is still limited understanding on how they affect drug response ^6^. Multiple efforts have been devoted to the large-scale *in vitro* screening of drugs across cell lines ^7–9^ that have proven useful to identify some genomic markers associated with drug response. However, molecular data alone has not proven sufficient to predict the efficacy ^10^ or toxicity ^11^ of a drug on an individual cell line in a reliable way. This predictability is likely to be even lower in patients, given the additional complexities when compared to cell lines.

Due to these limitations, genomics data has to be supplemented with other information in order to optimally guide the treatment for each patient, and systems for phenotypic stratification are urgently needed ^6,12^. The need for new approaches is even more acute for the application of drug combinations. Combinatorial targeted therapy has been shown to be a powerful tool to overcome drug resistance mechanisms, which can be due to tumor heterogeneity, clonal selection or adaptive feedback loops ^13^, and seems to be a particularly promising approach for treatment of pancreatic cancer ^6^. However, strategies to identify effective combinations are still in their infancy ^14^.

A powerful means to overcome these difficulties would be the ability to test the drug compound directly on patient samples. Despite recent progress towards functional testing of live patient tumor cells ^12,15^, the application of standard drug screening technologies is currently limited by the need of large numbers of cells ^15^. Therefore, large scale drug screening of patient tumors has been so far limited to blood tumors ^16^ (where a much larger amount of malignant cells is easily accessible) or requires some *ex vivo* culturing steps ^6^ (e.g. patient derived cell lines, PDX models and organoids ^17,18^) that require long times to grow the cells and can cause changes in the phenotype of the cells. Hence, there is, hence, no technology to perform drug screenings with the number of cells that can be obtained with biopsies from a solid tumor.

Microfluidic technology can in principle overcome screening restrictions based on limited starting material. Making use of tiny assay volumes, microfluidic systems have recently been applied successfully to the testing of a few individual drugs on primary tumour cells, tumour spheroids and tissue slices ^19–21^. However, these studies were based on single aqueous phase microfluidic systems which can process only small sample numbers (max ~96 including replicates, typically much less). A possible solution for further scale-up is the use of droplet microfluidics ^22^. In these systems aqueous droplets surrounded by oil serve as independent reaction vessels. Such systems have already been used for genetic assays of cancer cells ^23,24^, but they have not yet been applied to personalized phenotypic drug screens. This is probably due to the fact that encapsulation of different soluble drugs (rather than just different cells) into droplets or plugs (sequential aqueous segments in a microfluidic channel, capillary or piece of tubing; spaced out by oil or air) is technically still very challenging. Switching between multiple fluid sources requires robotic systems (e.g. sequentially aspirating samples from microtiter plates), which are rather slow, or complex microvalve technology (e.g. to switch between fluids injected in parallel) ^22^. While a combination of both approaches has been described ^25^; to date none of these plug-based systems has been used for phenotypic assays with human cells. This may be owing to incompatibilities, e.g. due to the fact that mammalian cells sediment and secrete proteins and metabolites causing the plugs to stick and break at the channel walls (so-called “wetting”; also promoted by growth factors in the media). In addition, a robust method for sample tracking is required. In previous studies ^25–27^ plugs were simply stored in a sequential fashion (within a microfluidic channel or a piece of tubing), wherein the common fusion or splitting of plugs causes “frameshifts” resulting in loss of information on sample composition.

Here, we present a platform that can overcome these limitations, enabling the screening of drug combinations on mammalian cells and patient biopsies in a plug format. Our approach requires significantly less cells compared to conventional, non-microfluidic formats and provides one to two orders of magnitude higher throughput (in terms of samples per experiment) than existing systems. Our platform, based on microfluidic Braille valves ^28,29^ and an external autosampler, can rapidly generate plugs containing cells, reagents of an apoptosis assay and systematic combinatorial drug cocktails. All workflows have been optimized to guarantee compatibility with live cells, and we furthermore implemented a fully scalable sample barcoding system. We demonstrate application of this platform for screening cell lines and, more importantly, patient samples *ex vivo.* Furthermore, pathway modeling of this data allows us to define apoptotic pathways in a patient-specific way. This should open the way for further translational efforts and clinical applications in the near future.

## Results

### Microfluidic platform for drug screening

We developed a plug-based microfluidic platform allowing us to rapidly screen a large number of combinations of chemical compounds on a very limited number of cells. Specifically, with a starting material of less than 1 million viable cells more than 1100 samples could be screened (10 chemical compounds in all the pairwise combinations, with about 20 plugs (i.e. replicates) per condition), each containing about 100 cells (**Fig. 1a**).

**Figure 1.**
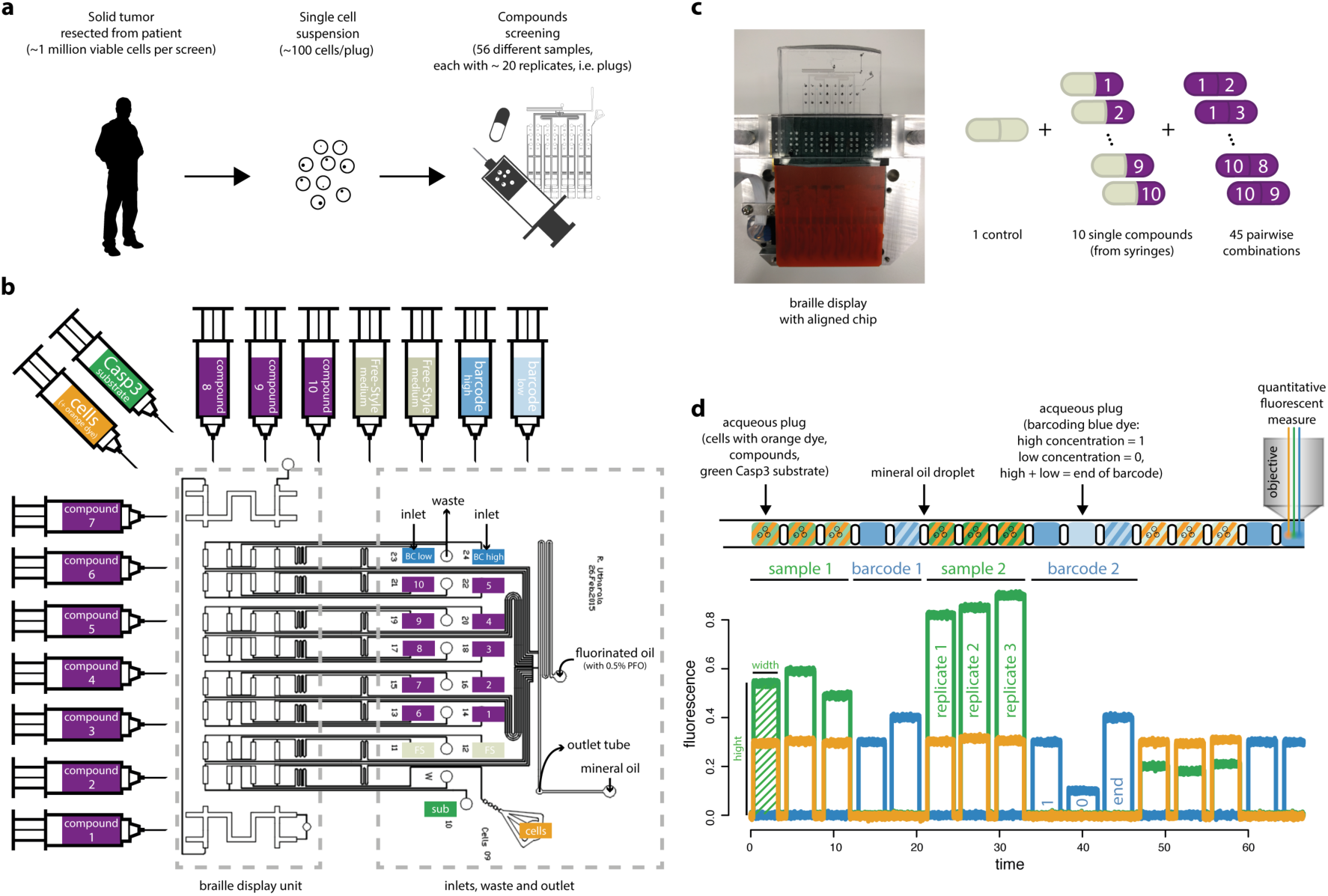
Microfluidics platform. (a) Workflow for patient samples. Functional drug screening on tumors resected from human patients: each resected tumor is dissociated to a single cell suspension, which is perturbed with different compounds using the microfluidics platform. (b) Chip design. 16 syringes with aqueous samples are connected to the inlets in the microfluidics chip via tubing (10 with compounds, 2 with medium to generate single drug and control samples, 2 for barcoding, 1 for the cell suspension, 1 with Caspase-3 substrate to detect apoptosis). Other 2 inlets in the microfluidics chip are used for carrier oil (FC-40) and mineral oil. The braille display unit is used to control the valves and regulate the flow coming from the aqueous phase syringes, resulting in different combinations. Plugs are collected in a tube connected to the outlet. (c) Experimental setup. The microfluidic chip is mounted on a Braille display and aligned with the pins. Single compounds and pairwise combinations are automatically generated. (d) Example of a plug sequence and corresponding readout. Multiple replicates (aqueous plugs) are produced for each sample. Each aqueous plug is generated by mixing the following components: cells (with orange fluorescent dye to verify the proper mixing of all components), caspase-3 substrate and one or two compounds. Sample plugs are followed by binary barcode plugs to encode the corresponding sample number (high concentration of blue fluorescent dye = 1, low concentration = 0, followed by an end of barcode signal).

Our platform is based on Braille valves ^28,29^ controlling individual fluid inlets of the microfluidic chip (**Fig. 1b-c**). All reagents (including the cell suspension, drugs and assay components) are permanently injected into the device and, depending on the valve configuration, sent either to the waste or to a droplet maker with a T-junction geometry. This approach allows rapid switching (~300 ms) between 16 liquid streams and, upon injection of fluorinated oil at the T-junction, the generation of combinatorial plugs at high throughput. If necessary, the chemical diversity can be increased further by connecting an autosampler to one of the inlets (sequentially loading compounds from microtiter plates; (**Supplementary Fig. 4**), but this procedure also requires a higher number of cells.

To avoid wetting issues we integrated an additional mineral oil inlet into our chip design, enabling the insertion of mineral oil droplets in between all (aqueous) sample plugs (**Supplementary Movie 3**). Such three-phase systems are particularly efficient in keeping samples separated ^30^, even under conditions that normally cause wetting. A further crucial factor for reducing wetting was the use of special, protein-free media. However, while a combination of these measures significantly improved reliability, plug integrity could still not be ensured for all samples. Hence a sample identification system that would still work if individual plugs break or fuse had to be implemented. Therefore, we introduced sequential barcodes to separate and identify samples of different composition, based on sequences of plugs with binary (high/low) concentrations of the blue fluorescent dye cascade blue. This way sample numbers can be encoded (e.g. high-low = binary number “10” = decimal number “2”). To demonstrate the power of this barcoding approach we converted a simplified EMBL logo (**Supplementary Fig. 3**) into a binary black and white image and translated all 2808 pixels into a sequence of plugs with two different fluorescence intensities. These barcodes were then detected using our laser spectroscopy setup (**Supplementary Fig. 1**) and converted back into the initial image, which did not show a single mistake. This clearly demonstrates the scalability and reliability of our strategy.

In each sample plug, cells are encapsulated together with one or two compounds and a rhodamine 110 (green-fluorescent dye) conjugated substrate of Caspase-3, which is an early marker of apoptosis ^31^. Furthermore Alexa fluor 594 (orange-fluorescent dye) was added to the cell suspension for verifying dilution by all assays reagents and monitoring correct operation of all valves. For each drug treatment multiple replicates were generated (see screening specific details below), followed by a fluorescent barcode, and stored sequentially in gas permeable tubing with a total length of 5m (having a capacity for about 3500 plugs in total). After overnight incubation, the readout was performed by flushing the samples through the detection module. In our case, we used an optical setup (**Supplementary Fig. 1**) with three different excitation lasers (375nm, 488nm and 561nm) and performed a multiplexed readout at three different wavelengths (450nm, fluorescence barcodes; 521nm, Caspase-3 activity; >580nm, orange marker dye to monitor mixing of reagents) using photomultiplier tubes (PMTs).

### Data extraction and quality assessment

In the fluorescence data, each plug corresponds to a peak in one or more channels (i.e. green, orange, blue), as shown in **Fig. 1d**. When processing the data, we first identified the peaks in the blue channel, representing the barcode, and we used them to separate and identify the peaks corresponding to each sample (Online Methods). Each sample is composed of multiple peaks (typically 12 replicates per run) with signals both in the green and in the orange channel. For each peak we considered two measures: height and width. The height of the peak is proportional to the measured fluorescence intensity: the intensity in the green channel represents the activation of Caspase-3 (thus apoptosis) while the intensity in the orange channel represents the concentration of the orange-fluorescent dye, which was added to the cell suspension. Since each plug was produced by mixing four components (cells, Caspase-3 substrate and two compounds/medium), the intensity in the orange channel allowed assessing the quality of this mixture and was used to discard samples/peaks with extreme values (i.e. outliers, see Online Methods). The width of the peak represents the length of the plug. Thin or wide peaks were discarded as they correspond to split or fused plugs respectively. Additionally, the first peak for each sample was typically discarded to avoid the effect of cross-contamination between samples.

### Combinatorial screening of pancreatic cancer cell lines

To optimize and validate our microfluidics platform we performed combinatorial screening of compounds on two pancreatic cancer cell lines with different genotype and phenotype ^32^: AsPC1 and BxPC3. Ten compounds were screened alone and in pairwise combinations (**Table 1**). We included in the screening: two drugs that are currently used in clinical chemotherapy as first line treatment for pancreatic cancer (Gemcitabine and Oxaliplatin), seven drugs that have specific kinase targets which play key roles in different pathways (i.e. IKK, MEK, JAK, PI3K, EGFR, AKT and PDPK1 inhibitors) and one cytokine (TNFα) which activates the extrinsic apoptosis pathway. The sequence of the samples is shown in **Fig. 2a** (data refer to BxPC3 cells, but sample sequence was the same for both cell lines): we screened 56 different samples, repeating the untreated control sample (i.e. only cells, Caspase-3 substrate and medium, without compounds) after every 10 samples and as first and last sample (total of 62 samples). For each sample we produced 12 replicates and considered the median value. The whole experiment was repeated six times (six runs) to verify the reproducibility of the measurements, resulting in about 6600 plugs in total including the barcode. In order to compare the different runs we computed the z-score for each (i.e. standardization by subtracting the mean and dividing by the standard deviation). Data (**Fig. 2a** for BxPC3 and **Supplementary Fig. 6** for AsPC1) show that control samples remained constant over time, proving that storing the cell suspension during plug production in the syringe on ice (protocol described in Online Methods) did not affect the viability of the cells. For further analysis the median values across the six runs were considered (**Fig. 2b,c**).

**Table 1.** List of screened compounds.

| **Compound name** | **Compound type** | **Putative target (effect)** |
| --- | --- | --- |
| ACHP | targeted | IKK (inhibition) |
| AZD6244 | targeted | MEK (inhibition) |
| Cyt387 | targeted | JAK (inhibition) |
| GDC0941 | targeted | PI3K (inhibition) |
| Gefitinib | targeted | EGFR (inhibition) |
| MK-2206 | targeted | AKT (inhibition) |
| PHT-427 | targeted | AKT, PDPK1 (inhibition) |
| Gemcitabine | cytotoxic | DNA replication (inhibition) |
| Oxaliplatin | cytotoxic | DNA replication (inhibition) |
| TNFα | cytokine | TNFR (activation) |

**Figure 2.**
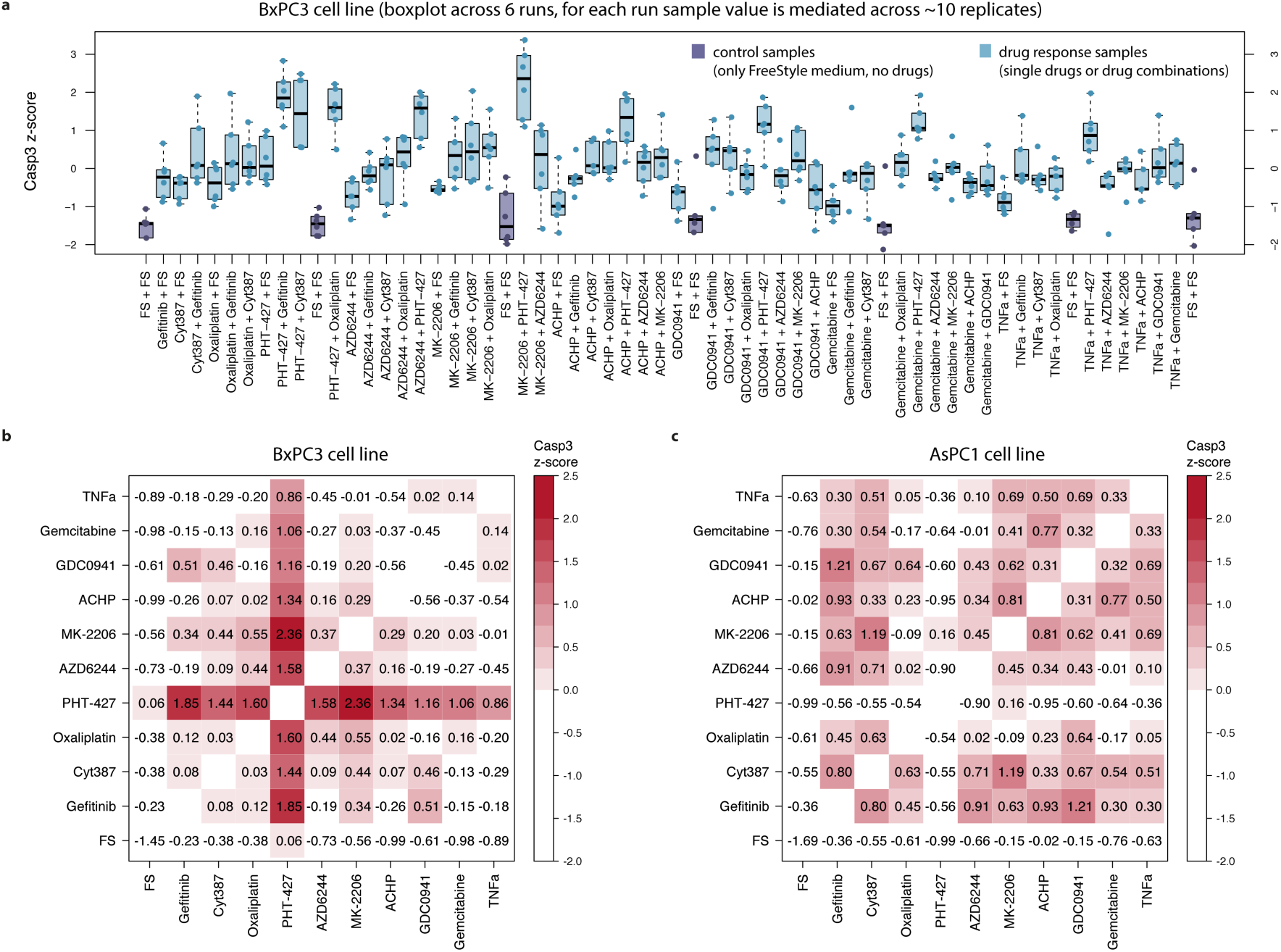
Microfluidic combinatorial drug screening in cell lines. (a) Boxplot of the sequence of samples across multiple replicates for BxPC3 cells (z-scores of Caspase-3 activity). (b) Heatmap representation of the same sample as in (a) (BxPC1 cell line) using median value across 6 replicates. Coloured red scale starting from z-score equal to 0 (i.e. median activation across all samples). (c) Heatmap representation for AsPC1 cells.

### Efficacious drug combinations and validation for screened cell lines

Combinatorial screening using the microfluidics platform suggested potentially interesting drug combinations (**Fig. 2b,c**). Some combinations showed strong effects in both cell lines (e.g. GDC0941 and Cyt387, z-score = 0.46 for BxPC3 and 0.67 for AsPC1), while, more interestingly, some stronger effects were shown to be specific for each cell line. In particular, PHT-427 induced apoptosis in BxPC3 cells in combination with multiple drugs (z-score range = [0.86, 2.36]), while it showed little or no effect on AsPC1 cells (z-score range = [−0.95,0.16]). The strongest effect for BxPC3 was measured in response to combinatorial treatment with PHT-427 (AKT and PDPK1 inhibitor acting on the PH domain) and MK-2206 (allosteric inhibitor of AKT). We compared the combinatorial effect on BxPC3 with the corresponding single drug treatment (**Fig. 3a**); either single treatment showed an effect that was not significantly higher than zero (p-value 0.21 and 1 for PHT-427 and MK-2206 respectively, one-tailed t-test), while the combinatorial treatment showed a strong and significant effect (effect size = 2.24, Cohen’s d; p-value = 0.001, one-tailed t-test). On the contrary, no significant effect was shown for AsPC1 for PHT-427, MK-2206, or their combination (p-value = 1, 0.68 and 0.28 respectively, one-tailed t-test), suggesting that the efficacy of the drug combination on BxPC3 is not caused by a general toxicity of the drugs when administered in combination, but rather by a cell line specific effect. Similarly, a promising treatment specific for AsPC1 cells is the combinations of Gefitinib (EGFR inhibitor) with ACHP (IKK inhibitor), which showed a strong efficacy for AsPC1 (effect size = 1.34, Cohen’s d; p-value = 0.01, one-tailed t-test) but not for BxPC3 (p-value = 0.83, one-tailed t-test). For both examples, results were validated in subsequent tissue culture experiments (**Fig. 3b,d**), confirming the behavior observed in the microfluidic system.

**Figure 3.**
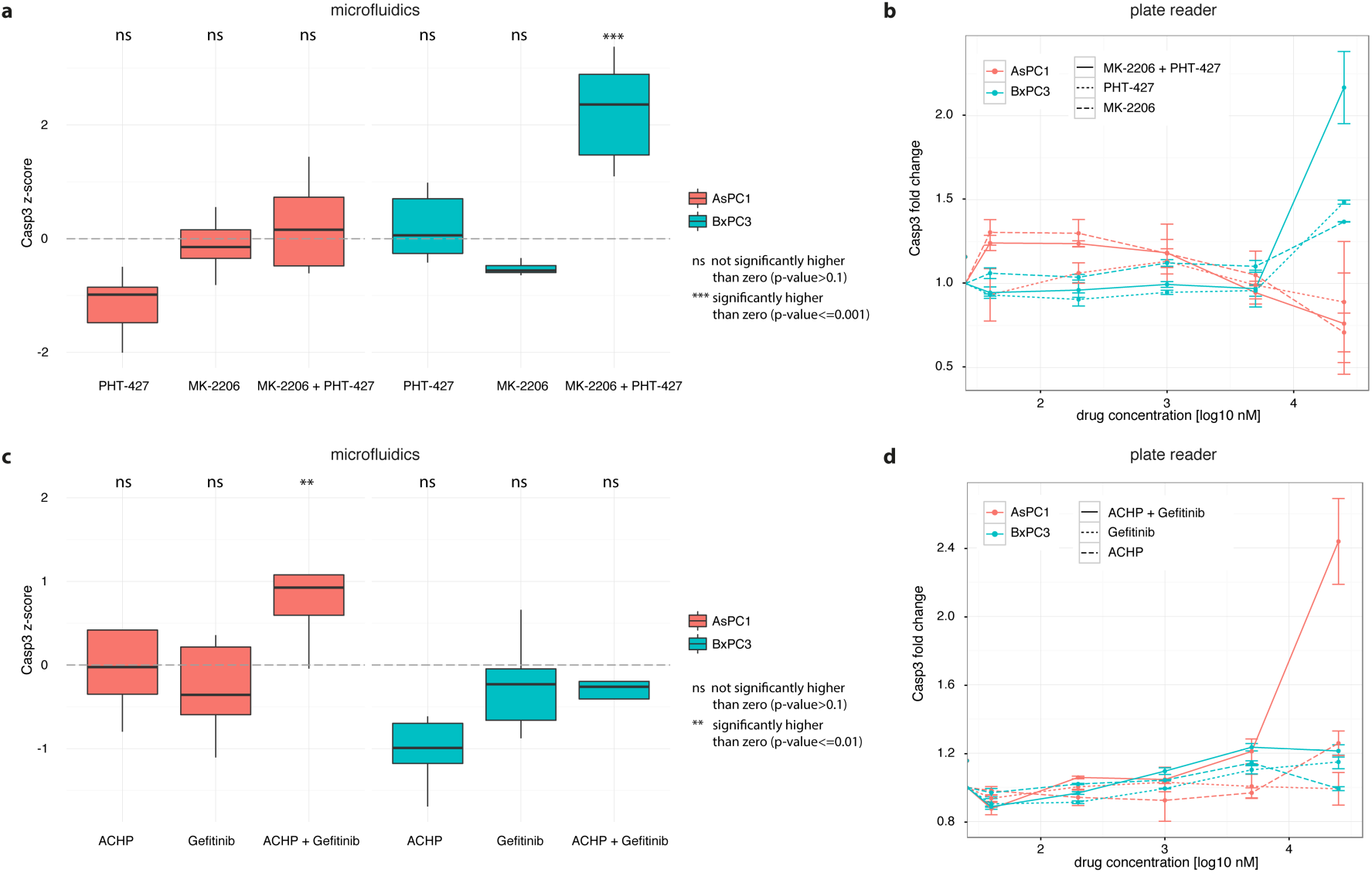
Validation of cell line specific drug combinations in tissue culture experiments. (a) BxPC3 show a strong activation in the microfluidic system when treated with the combination of MK-2206 and PHT-427, which is not seen when treated with the single drugs. No strong activation is shown for AsPC1 treated with the same drugs. (b) Same behaviour is confirmed in 96-well plates where the drug combination shows a much higher signal than the two single drug samples for BxPC3 cells, while it stays at the basal level for AsPC1 cells. (c,d) Similarly, the combination of ACHP and Gefitinib is potent in AsPC1 cells but not in BxPC3 cells, both in microfluidic plugs (c) and in a 96-well plate format (d).

### Combinatorial screening of resected patient pancreatic tumors

The same set of ten compounds used on cell lines was then applied to screen tumor samples resected from 4 pancreatic cancer patients. For each patient sample, the solid tumor was dissociated to create a single cell suspension that was then used for the screening (protocol in Online Methods). Similar to the pipeline used to screen the cell lines, a total of 62 samples were produced, with 12 replicates each (each plug with about 100 live cells). The whole sample sequence was repeated two or three times (different runs) depending on the amount of viable cells that were obtained from the patient biopsy. As in the case of the cell lines, unreliable peaks/samples were discarded based on the intensity in the orange channel and on the length of the peaks. Only samples passing the quality assessment in at least two runs and with consistent values across runs were considered for further analysis (boxplot with data for each run shown in **Supplementary Fig. 7**, more details in Online Methods).

As shown in **Fig. 4a–d,** some drug combinations showed promising results in more than one patient, but no combination was effective for all patients. In particular, the combination of PHT-427 and ACHP was very effective in three patients (patients #1, #2, #4 with z-score equal to 1.03, 1.39 and 1.64 respectively), while showing no apoptosis induction in one patient (patient #3, z-score equal −0.44). Other combinations, i.e. Cyt387 and PHT-427, PHT-427 and AZD6244, PHT-427 and GDC0941, showed strong response (i.e. z-score higher than 0.9) for two patients while showing low or no apoptosis induction in the remaining patients. Interestingly the best combination (highest z-score) was different for each patient: PHT-427 and AZD6244 for patient #1, PHT-425 and ACHP for patient #2, MK-2206 and GDC0941 for patient #3, MK-2206 and ACHP for patient #4. This clearly illustrates the need for personalized approaches.

**Figure 4.**
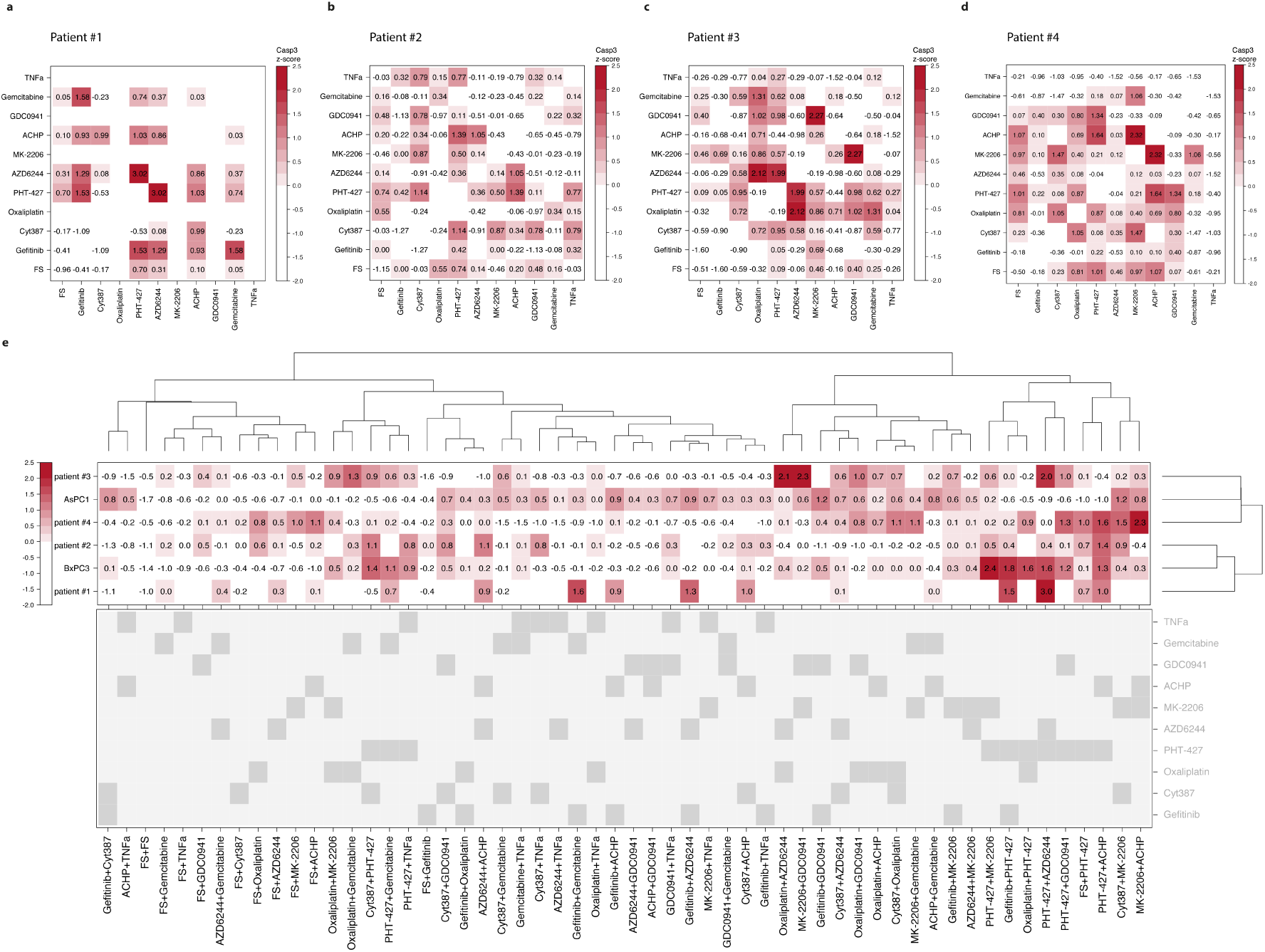
Patient samples and comparison with cell lines. (a-d) The heatmaps show the median z-score of Caspase-3 activity for each patient sample for which data could be obtained successfully. No value is reported for samples that were not measured (two of the 10 drugs were not screened for patient 1) or had technical issues (e.g. samples showing unexpected dilutions of the orange marker dye added to the cell suspension) or that had inconsistent values between runs. Coloured red scale starts from z-score equal to 0 (i.e. median activation across all samples). (e) Data clustered by samples and by patient/cell line.

Gemcitabine and Oxaliplatin, the current first-line treatments for pancreatic cancer showed increased apoptosis induction when administered in combination with each other (in patient #3, z-score equal 1.31) or with other drugs. In particular, Gemcitabine strongly induced apoptosis in combination with Gefitinib for patient #1 (z-score equal 1.58) and with MK-2206 for patient #4 (z-score equal 1.06), while Oxaliplatin is effective in combination with AZD6244 or GDC0941 for patient #3 (z-score equal 2.12 and 1.02 respectively) and with Cyt387 for patient #4 (z-score equal 1.05).

### Comparison between patient samples and cell lines

Clustering the screening data by individuals (patient/cell line) and by samples (**Fig 4e**) showed how most of the single compounds and some combinations, e.g. AZD6244 and TNFα, failed to show strong efficacy on any individuals. Other treatments displayed high efficacy both on patient samples and on cell lines (e.g. combination of PHT-427 and ACHP). Importantly, none of the treatments is effective across all individuals, suggesting that they are not widely toxic, but rather patient specific. This further demonstrates the need for individualized treatments.

### Network-based interpretation

In order to better understand the mechanism of action of the screened compounds, we derived a logic model of pathways involved in apoptosis (**Fig. 5a**) from literature ^33–37^, describing both intrinsic (mediated by the mitochondria, named Mito in the model) and extrinsic (mediated by Tumor Necrosis Factor Receptors TNFR) apoptotic signals, including nodes encoding for both anti- and pro-apoptotic effects. We incorporated in the model all nodes perturbed by specific compounds in our screening such as targeted drugs (kinase specific inhibitors) and cytokine TNFα. The effect of chemotherapeutic (DNA damaging) drugs could not be included in the model since they inhibit DNA replication rather than acting on specific signaling nodes. To encode for the different mechanisms of action of MK-2206 and PHT-427 on AKT (allosteric and PH domain inhibitors respectively), they were modeled as acting on two different nodes (AktM and AktP respectively), both needed for the activation of AKT.

**Figure 5.**
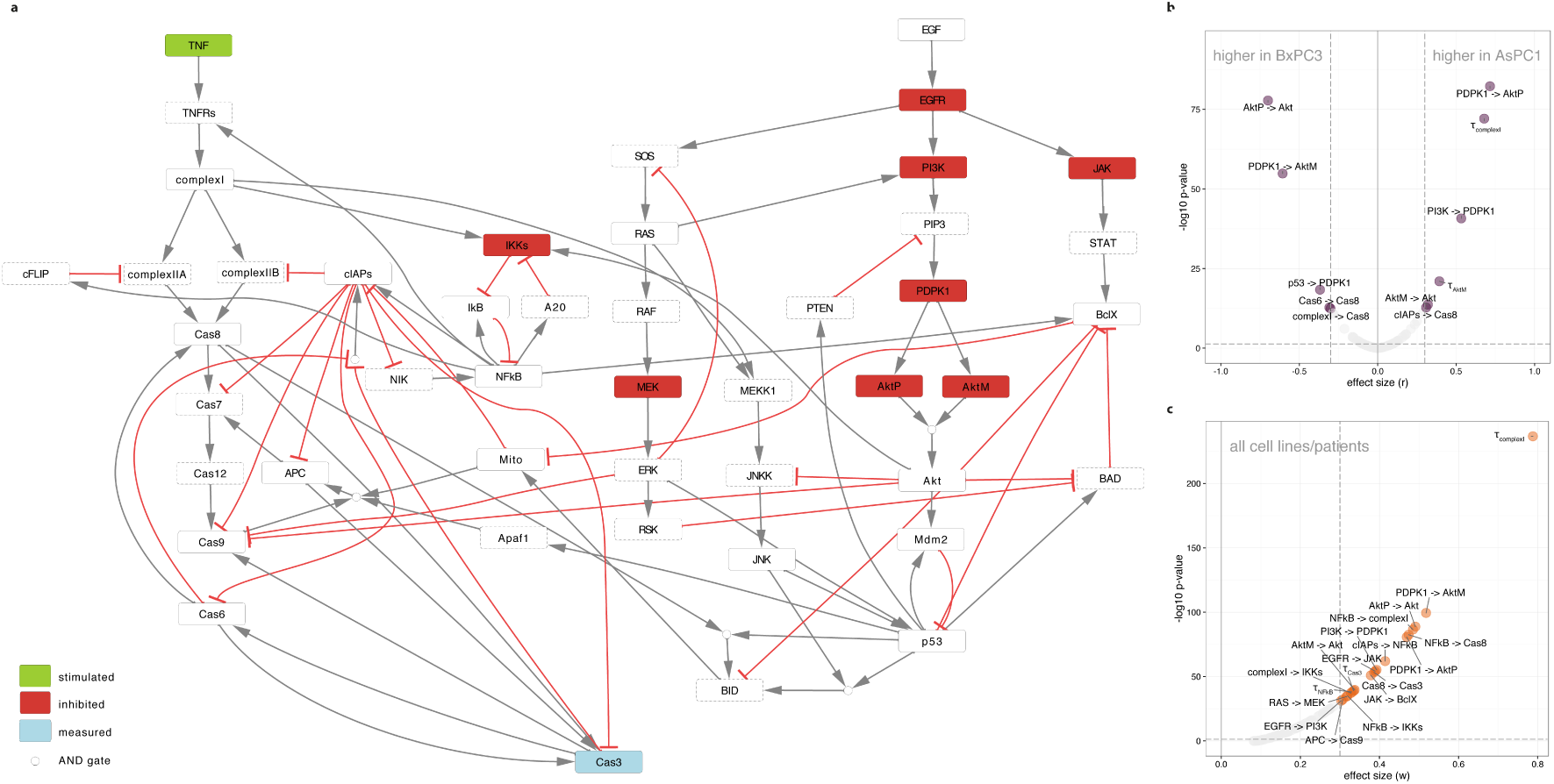
Cell line/patient specific models of apoptosis pathway. (a) General structure of the logic model of intrinsic and extrinsic apoptosis pathways. Nodes perturbed in the combinatorial drug screening are shown in green if stimulated (cytokine TNFα) and in red if inhibited; measured node (Caspase-3, shortened to Cas3 in the figure) is in blue; dots represent AND gates. Dotted nodes are compressed before estimating cell line/patient specific parameters. Edges are shown in black for activation and in red for inhibition. (b) Volcano plot representing the results of the comparison of the bootstrap distribution of the parameters for AsPC1 and BxPC3 cells. (c) Volcano plot representing the results of comparing the bootstrap distribution of the parameters across all cell lines/patients.

Cell line and patient specific models were built using a logic ordinary differential equations formalism ^38^ as implemented in CellNOpt ^39^. The starting general model structure was fitted to the experimental data for each cell line and patient (see Online Methods). Estimated parameters are one life-time parameter for each species (e.g τ_EGFR_ for node representing EGFR) and a regulation parameter for each interaction (e.g. EGFR → JAK). For each cell line/patient a bootstrap distribution was obtained for each parameter (resampling experimental data with replacement 300 times).

In order to identify the pathway differences that might underlie the differential response to the drugs in our two tested cell lines, we compared the distributions of the parameters using Wilcoxon rank sum test for assessing the significance (p-values Bonferroni corrected for multiple hypothesis testing > 0.05) and *r* effect size (medium-high effect size > 0.3, see Online Methods); results are shown in (**Fig. 5b**). Similarly, distributions across all cell lines/patients were compared using Kruskal-Wallis rank sum test (one-way analysis of variance, p-values Bonferroni corrected > 0.05, see Online Methods; **Fig. 5c**). The main differences between AsPC1 and BxPC3 pathway models involve the extrinsic pathway and the Akt pathway: AsPC1 cells have higher positive regulator parameter values (which correspond to a slower response) for parameters involved in the regulation of Akt initiated by PI3K (i.e. PI3K → PDPK1), while BxPC3 cells have higher values for the negative feedback to Akt, mediated by PTEN (i.e. p53->PDPK1). Other significantly different parameters involve nodes used to model the different mechanism of action of our two Akt inhibitors (i.e. AktM and AktP) and are strongly influenced by the differences that the two cell lines show especially in response to PHT-427. Interestingly, it has been previously shown that alterations in the activity of the AKT pathway are quite common in pancreatic adenocarcinoma due to mutation and epigenetic alterations ^40,41^.

## Discussion

We describe here a microfluidic platform enabling the fast screening of many drug combinations in a personalized approach. The very small assay volumes (~100 cells per 0.5 μL plug) allowed us to perform comprehensive screens directly on patient biopsies. Importantly, these can be done without the need for any intermediate cultivation steps, which might introduce significant cellular changes and/or the selection of individual clones. Hence our approach opens the way for comprehensive screens that have so far been restricted to blood tumours, for which patient-derived cancer cells are available in large quantities ^16^. Furthermore, the low volume and the high level of automation allow such screens to be rapidly performed at low cost: Within 48h after tumour resection or biopsy, and with consumable costs of about US$ 150, which is considerably lower than routine procedures such as MRT scans or surgical interventions. Therefore, we envision rapid translation of this technology into clinical application.

Functional combinatorial screening has great potential in predicting personalized therapy as revealed by the data presented in this paper. Our cell line screening recapitulates the lower sensitivity of KRAS mutants (AsPC1, vs wild type BxPC1) to PHT-427 ^42^, (Fig. 1 b-c, for the single drug treatment as well as across combinatorial treatments involving PHT-427). In addition, we could suggest and validate a novel and particularly strong and specific drug combination for BxPC3 when treated with PHT-427 and MK-2206. These two compounds are both AKT inhibitors, although they act through different sites, and PHT-427 additionally inhibits PDKP1 ^42^. Subsequently, similar efficacious specific combinations were suggested for each patient sample. Interestingly, no combination showed strong efficacy across all cell lines/patients suggesting that the tested treatment options are not just generally toxic but rather cell line/patient specific. This inter-patient heterogeneity in response to treatment highlights the importance of our approach to personalised medicine and is in agreement with previous findings ^6^, where sensitivity profiles were shown to be particularly patient specific for pancreatic cancer.

A network-based perspective can be informative when investigating the best combinatorial therapies ^43^. For example, mapping the effect of the kinase specific inhibitors on the signaling pathways can help to prevent the mechanisms of drug resistance by suggesting combinatorial therapies that act on parallel pathways. An interesting example is the combination of PHT-427 (AKT inhibitor) and AZD6244 (MEK inhibitor), which was effective (z-score > 0.4) in three out of the four tested patient samples. These drugs in fact block two important parallel pathways which are the MEK-ERK and the PI3K-AKT pathways. Although, as far as we know, synergistic combinations targeting these pathways have been previously studied for pancreatic cancer, especially in the context of KRAS mutants (>90% of pancreatic cancer patients are KRAS mutants)^44^, this is the first evidence of potential synergistic effect between PHT-427 and AZD6244. The efficacy of this treatment also in BxPC3 cells (KRAS wild type) might suggest a more general applicability.

Perturbation experiments with targeted drugs (e.g. kinase-specific inhibitors) can also be used to infer the cell line or patient specific pathways ^45^. Signal transduction models can also be linked to cell viability/apoptosis ^46^ in order to predict drug sensitivity. We used data from our perturbation experiments and prior knowledge on the signaling pathways impinging on apoptosis to build cell line and patient specific models which were then used to unveil potential biological reason for the differential response. Models could, in principle, be in the future also used to predict new potentially interesting targets.

Compared to other personalized approaches our approach has specific advantages: While organoids and xenografts are particularly well-suited for mimicking three-dimensional tumour architecture and the *in vivo* microenvironment, the use of individualized tumour cells as shown here facilitates rapid, massively parallelized assays at low cost. This could be exploited further by implementing single-cell droplet assays ^22,47,48^, which require fewer cells and also allow to take intratumoral heterogeneity into account. In addition to deriving an averaged readout over 50 single-cell replicates per treatment option, one could quantitatively determine the number of non-responding cells for each drug cocktail. Such data would probably be very valuable to overcome therapy resistances.

Apart from using it for personalized cancer therapies, our microfluidic platform should be of interest for further applications, such as the screening of cocktails for targeted stem cell differentiation ^49,50^ or combinatorial chemistry ^51^. We have shown for the first time how an affordable and robust Braille display (< US$ 1000) can be used to combine microfluidic valve and droplet technology. This way droplets of different chemical composition can be rapidly generated on demand, without the need for multi-layer channel networks and/or the use of expensive multi-channel pressure controllers. Also of note is the fact that we used only 32 of the 64 available Braille pins, which leaves further room for scaling-up, especially as there are even bigger Braille displays with more than 300 pins. In addition, the possibility of stopping the flow in individual channels of a network should facilitate the construction of cheap and versatile multi-way cell- and droplet-sorters.

## Author contributions

F.E. designed and performed experiments, performed the computational analysis under the supervision of J.S.R. and shaped the overall study design. R.U. designed and performed experiments and developed the microfluidic platform under the supervision of C.A.M. D.M. performed experiments. U.P.N. supervised the surgical specimen collection and provided clinical patient data. T.C. provided clinical samples and contributed to the design of the study. J.S.R. and C.A.M. conceived the project, designed experiments and supervised the study. F.E., J.S.R. and C.A.M. wrote the manuscript. F.E., R.U., D.M., T.C., J.S.R., C.A.M. interpreted the results and contributed to manuscript development. All authors approved the final manuscript.

## Acknowledgments

We gratefully acknowledge all patients who kindly gave their consent for research use of the biopsies. We thank Ilka Sauer, Anne Esser and Ellen Krott for highly appreciated technical assistance, EMBL electronic workshop for building Braille controllers and implementing a LabVIEW command list, EMBL mechanical workshop for building customized parts for the Braille display, Felix Krüger and John P. Overington for help selecting kinase specific inhibitors, Diana Panayotova Dimitrova and Maria Feoktistova for constructive discussion about apoptosis pathway, Jan Korbel and Denes Turei for critical reading of the manuscript. F.E. thanks European Molecular Biology Laboratory Interdisciplinary Post-Docs (EMBL EIPOD) and Marie Curie Actions (COFUND) for funding.

## Online Methods

### Microfluidic setup

Microfluidic chips were fabricated using standard soft-lithography methods. In brief, molds were fabricated on 4-inch silicon wafers (Siltronix) using AZ-40XT positive photoresist (Microchemicals) according to the manufacturer's instructions. Patterning was achieved by projecting 25400 dpi photomasks (Selba) onto the photoresist using light with a wavelength of 375 nm (Suess MicroTec MJB3 Mask Aligner). All channels had a height of ~50μ.m and widths ranging from 50 μm - 400 μm (400 μm for all valve sections). PDMS chips were manufactured using elastomer and curing agent in a ratio of 1:10 (Sylgard 186 elastomer kit, Dow Corning Inc) cured overnight at 65°C. To allow for valve actuation (compression of the valve sections by the Braille pins) the PDMS chips were not bonded to glass, but rather to a thin elastic PDMS membrane. This membrane was prepared using elastomer and curing agent in a 1:20 ratio, poured on a transparency sheet and spin coated at 700 rpm (Laurell WS 650) before overnight curing. Bonding was performed in a Diener Femto Plasma Oven. Connections to inlets and outlets were punched using 0.75mm Harris Unicore Biopsy punches. Before use, chips were flushed with Aquapel (PPG industries) from the outlet up to the T-junction to render the channel surface hydrophobic.

For screening, the Braille valve chip (**Fig. 1B**) was mounted onto an SC-9 Braille display (KGS Corporation, Japan) using an in-house holder as shown in **Fig. 1C**. This holder includes a Plexiglas bar with built-in screws to push the PDMS chip onto the Braille pins. The design of the chip ensures that all fluid connections (inlets and outlets) are outside the area covered by the Plexiglas bar. Movement of the Braille pins was controlled by an in-house LabVIEW programme, enabling to actuate all individual pins according to a pre-defined sequence (corresponding to systematic sample combinations and barcodes). Barcodes were generated using binary concentrations of Cascade Blue (16 μM and 48 μM for low and high respectively).

For all experiments the aqueous fluids were injected at a flow rate of 500 μl/h using Harvard Apparatus Syringe pumps. FC-40 containing 0.5% perfluoro-octonoal (PFO, ABCR) and mineral oil were injected at rates of 200 μl/h and 175 μl/h. Resulting aqueous plugs were 1.5 mm to 2 mm in length and 424 nL to 560 nL in volume.

Subsequent to incubation, the fluorescence of all samples was determined inside PTFE tubing (Adtech) with an inner diameter of 600 μm using the optical setup shown in **Supplementary Fig. 1**. The resulting sample data was analyzed using custom R-scripts (see paragraph “data processing” for details).

### Setup of the fluidic system, choice of additives and oils

We used fluorinated oil without stabilizing surfactant as a carrier phase (only 0.5% of the anti-wetting agent PFO was added), which turned out to have two major advantages: upon reaching the outlet, small droplets generated at the T-junction fused and formed larger plugs that completely filled the collection tubing, thus allowing to incubate all samples in a sequential order. Furthermore the lack of surfactant prevented the formation of micelles, which can cause the exchange of reagents between droplets ^52^. To increase stability of the arrays, plugs were furthermore spaced out using mineral oil (Sigma).

### Integration of an autosampler

To allow for upscaling of the screens, we integrated an autosampler into our microfluidic platform ^27^. One of the inlets of the microfluidics chip was connected to a Dionex 3000SL Autosampler, aspirating samples from 96-well plates and injecting them sequentially into a target tubing connected to an external Harvard Apparatus Syringe pump. While the resulting throughput for loading compounds from different wells is rather low (~90 sec per reagent), each of them can be mixed with all of the drugs injected directly into the Braille valve chip. Therefore, the maximal throughput in terms of pairwise combinations is much higher (e.g. 9 sec per combination when mixing with 10 further reagents directly connected to the Braille display chip). However, one effect had to be taken into account: the concentration of compounds coming from the autosampler varied due to dispersion of the samples in the miscible carrier phase (the buffer used in the robotic system) ^53^. To overcome this effect, we implemented a feedback loop between the autosampler and the Braille valves: The relay signal of the autosampler was used to send the beginning and end of each sample from the microtiter plates to the waste, using the two Braille valves controlling the relevant inlet on the microfluidic chip. This process allowed to overcome sample dispersion and is illustrated in **Supplementary Fig. 4**. This way only the centre part of each sample segment, showing constant concentration, was mixed with the drugs injected directly into the Braille display chip. To verify this procedure we mixed fluorophores stored in a 96-well plate with fluorophores injected directly into the autosampler and measured the resulting fluorescence signal of the droplets (**Supplementary Fig. 5**). All combinations showed the expected signal intensities, confirming the feasibility of the approach if a sufficient amount of cells is available for screening.

### Single cell suspension from pancreatic primary tumors

Primary pancreatic tumors were obtained from routine resections from patients who signed an informed consent approved by the Research Ethics Committee of the Medical Faculty of the RWTH Aachen University (EK 206/09). The project was also approved by the EMBL Bioethics Internal Advisory Committee. A viable single cell suspension was prepared from the fresh tumors and directly used in an experiment within the next few hours. Tumors were first mechanically dissociated (~2–3 mm pieces) and digested for 1.5 hour at 37° C in 5ml of prewarmed digestion media consisting of 1 mg/ml collagenase (Sigma) solution in DMEM/F12 (Gibco, Life Technologies). Solution was pipetted every 30 min to facilitate dissociation, diluted in 25 ml of buffer (PSB) to stop the reaction, and centrifuged at 250 g for 5 min. Supernatant was then removed and 2 ml of 0.05% trypsin-EDTA (Gibco, Life Technologies) were added and the solution was incubated for 5 more minutes at 37° C. Subsequently 15 ml of DMEM supplemented with 10% fetal bovine serum were added and the solution was centrifuged again for 5 min at 250 g. After removing the supernatant, the pellet was resuspended in FreeStyle medium (ThermoFisher).

### Preparation of cells for microfluidic experiments

Cultures were trypsinized to detach cells, harvested and washed with PBS. The same procedure was used to prepare the single cell suspension from either primary tumor or cell lines for the microfluidic experiment. Cells were suspended in FreeStyle medium with the addition of 1 mg/ml Xanthan Gum (Sigma), for density matching, and 2 μl/ml of 10% Pluronic (Sigma), to reduce cell attachment and formation of clumps. Cells were filtered using a 40 μm cell strainer and diluted to a final concentration of 8×10^5^ viable cells/ml (counted with trypan blue exclusion method using a BioRad cell counter). 15 μg/ml of the orange fluorescent dye Alexa fluor 594 (ThermoFisher, #A33082) were added to the cell suspension in order to verify the proper mixture of the components in the plug. Cell suspension was then pipetted in a 5 ml syringe with the tubing directly connected to the syringe (no needle to avoid clogging) using PDMS and UV glue, and with a magnetic stir bar inside the syringe. For the duration of the experiment cells were maintained at low temperature using ice and constantly stirred.

### Preparation of drugs and Caspase-3 substrate

Cyt387 (#S2219), PHT-427 (#S1556), MK-2206 (#S1078), GDC0941 (#S1065), Gefitinib (#S1025), Oxaliplatin (#S1224), AZD6244 (#S1008) and Gemcitabine (#S1149) were purchased from Selleckchem. ACHP (#4547) was purchased from Tocris. All compounds were diluted in DMSO to a 20 mM stock solution. When preparing the syringes for the microfluidic experiment, compounds were further diluted in FreeStyle medium to a final concentration of 5 μM in the plugs. Tumor Necrosis Factor-α (TNF) (#PHC3015) was purchased from Life Technologies and diluted according to the manufacturer's instruction to a 10 μg/ml stock solution. It was further diluted in FreeStyle medium for microfluidic experiments to a final concentration of 5 ng/ml. For validation experiments in 96 well plates, compounds were instead prepared in 5 consecutive 5-fold dilutions (25 μM, 5 uM, 1 uM, 0.2 uM, 0.004 uM).

The caspase-3 substrate (Z-DEVD)2-R110 was purchased from Biomol (#ABD-13430). 3 ml of substrate working solution were prepared by adding: 2400 [μl 5X

Reaction buffer (20 mL of 50 mM PIPES, pH 7.4, 10 mM EDTA, 0.5% CHAPS), 60 μl DTT (1M), 540 μl dH2O and 44 μl Z-DEVD-R110 substrate (5mM).

### Data processing

Data were acquired using an in-house LabVIEW program allowing the detection of three channels (i.e. blue = fluorescence barcodes, green = Caspase-3 activity and orange marker dye to monitor mixing of reagents) as exemplified in **Fig 1d**. Peaks in each of the three channels were identified by defining an empirical threshold both on the intensity and the duration of the measured signal in order to distinguish real peaks from background noise. Peaks in the blue channel (corresponding to the barcode plugs) were detected and used to separate the different samples consisting of sequences of peaks (replicates) in the green channel. Samples with multiple peaks having either very low or very high orange signals were manually discarded (can be due to temporary cells clogging). Additionally, we also discarded peaks showing very high and very low width (based on empirical thresholds) in order to remove peaks corresponding to fused or split plugs respectively. After these steps, we considered the distribution of the intensity of the orange peaks across all samples and discarded the extreme values (i.e. the outliers). Where Q1 is the 25^th^ percentile, Q3 is the 75^th^ percentile and IQR is the interquantile range (Q3-Q1), outliers were defined as values lower than Q1 - 1.5*IQR, or higher than Q3+1.5*IQR. These strict rules were applied to guarantee higher quality of the data used for the analysis described in the paper. Code used to process the data is available as an R package in GitHub (https://github.com/eduati/BraDiPluS).

### Apoptosis pathways model

The logic model shown in **Fig. 5** was derived by manual literature curation starting from the model described by Mai and colleagues ^33^ and integrating additional information in order to include all nodes perturbed in our experiments and to well describe pathway cross-talks. We modeled both the intrinsic (mediated by the mitochondria) and the extrinsic (mediated by death receptor TNFR) apoptosis signal including nodes encoding both anti- and pro-apoptotic effects. Binding of TNFα to TNFR activates the extrinsic pathway mediated by Caspase-8 (Cas8 in **Fig. 5**) activation of Caspase-3 (Cas3 in **Fig. 5**). The two distinct Caspase-8 activation pathways ^54^ are represented by the cascade involving complex I (composed of RIPK1, TRADD, TRAF2), which induces the formation of two different Caspase-8 activation complexes: complex IIA (TRADD, RIPK1, FADD, Pro-caspase 8) and complex IIB (RIPK1, TRADD, FADD, Pro-Caspase 8, cFLIP) that can be inhibited by cFLIP and cIAPs respectively. For simplicity, Caspase-8 is modeled as a separate node (Cas8) regulated by the two complexes. TNFα can also regulate the intrinsic pathway through the activation of NFkB (anti-apoptotic node) by removal of its inhibitor IkB. The activation of the intrinsic pathway is executed by the mitochondria through the release of SMACs (second mitochondria-derived activator of caspases) and Cytochrome c. The former deactivates IAPs, which are anti-apoptotic proteins, the latter binds to Apaf1 (Apoptotic protease activating factor-1) and pro-caspase9 which is converted to its active form of Caspase-9 (Cas9 in **Fig. 5**) and in turn activates Caspase-3 (Cas3 in **Fig. 5**). Both Akt and ERK have an anti-apoptotic effect by phosphorylating BAD ^36^ and thus unbinding it from BclX and this can be modelled as an OR gate ^55^. We also included the pro-apoptotic effect of ERK as regulator of p53 ^37^. Additional cross-talks between the pathways (i.e. RAS → MEKK1 and RAS → PI3K) were included as described in ^35^. The resulting logic model described in **Fig. 5** was considered as a Prior Knowledge Network (PKN) and was then optimised based on the experimental data for each patient/cell line in order to generate patient/cell line specific models. We used a formalism based on logic ordinary differential equations ^38^ where ordinary differential equations are derived from logic models using a continuous update function *B_i_* for each species *x_i_*, which can assume continuous values {0,1}:

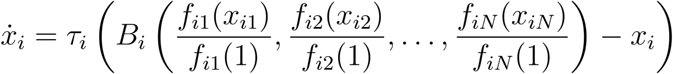

Where *x_i_*_1_,*x_i_*_2_… *x_iN_* are the N regulators of *x_i_* and each regulation is defined by Hill functions *f_ij_* with parameters *n_ij_* and *k_ij_*

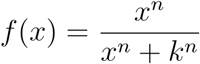

Optimization was performed using CNRode add-on of the CellNOptR package ^39^: parameters *τ* and *k* were estimated, while parameters *n* were fixed to 3. The notation used in the paper and in **Fig. 5** is τ_A_ for the life-time parameter τ for species A and A → B for the regulation parameter *k_AB_* The PKN was first compressed to reduce the complexity of the model by removing nodes that do not affect the logic consistency of the model ^56^. Compressed nodes are shown with dashed borders in **Fig. 5**.

Optimization for each patient/cell line was repeated 300 times with bootstrap (sampling training data with replacement) to obtain a distribution for each parameter. To compare the distributions of the parameters between the two cell lines we used Wilcoxon Rank sum test (as implemented in the R package ‘coin’), using 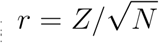 _as_ effect size, where *Z* is the statistics from the test and *N* is the number of observations. Effect size > 0.3 is considered as medium-large effect. To compare the distributions of all patients/cell lines we used Kruskal-Wallis rank sum test (which is a one-way analysis of variance on ranks) to test if observations derive from the same distribution for all groups, i.e. patient/cell line (null hypothesis rejected if different for at least one group). Effect size *w* is computed as 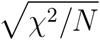 where χ^2^ is the statistics from the test and *N* is the number of observations. Effect size >0.3 is considered as medium-large effect. Rank type test were preferred over parametric tests because they are highly robust against nonnormality.

